# How best to co-deploy insecticides to minimise selection for resistance

**DOI:** 10.1101/2023.04.15.536881

**Authors:** Sam Jones, Andy South, Jason Richardson, Sarah Rees, Ellie Sherrard-Smith, Graham Small, Derric Nimmo, Ian Hastings

## Abstract

Insecticides are widely used to control the insects that spread human infectious diseases, in particular falciparum malaria. This widespread use has driven insecticide resistance (IR) to high levels that may threaten the effectiveness of future control programmes. There is interest in identifying deployment methods that alleviate the pressures driving IR and we investigate three. Mixtures are, as already known, highly effective in slowing IR providing their effectiveness (ability to kill fully sensitive insects) remain close to 100%. Mixtures may be expensive and/or operationally difficult so two alternatives to mixtures were investigated. Panels, where different insecticides are physically closely adjacent, for examples, different panels on the same bednet; mosquitoes may therefore encounter both insecticides in the same foraging cycle. Micro-mosaics where different insecticides are deployed in slightly wider geographic proximity, for example in adjacent dwellings. The mosquitoes are unlikely to encountered both insecticides in the same foraging cycle but may encounter different insecticides in subsequent foraging. It is hoped that panels and/or micro-mosaics may, by allowing individual mosquitoes to potentially encounter both insecticides, be effective, lower-cost alternatives to mixtures. Our results suggest this is unlikely to be the case. When insecticides are fully effective then mixtures remain clearly the best strategy. As effectiveness falls then all three strategies are roughly equal. The operational decision of what deployment methods to use depends on how confident we are that insecticides will have high effectiveness that will be maintained in realistic field conditions post-deployment.

## Introduction

Insecticides have been widely, and successfully, deployed to control mosquitoes that spread human disease; perhaps inevitably, this has driven widespread insecticide resistance. In particular, there is considerable concern that resistance to pyrethroids may undermine these control efforts (see, for example, [1–5] for recent reviews). This led the WHO to recommend that insecticides be deployed in ways that minimise selection for resistance ([6]; see further discussion in [7]). These deployment methods have become known as insecticide resistance management (IRM) strategies. Three main sources of evidence were used to identify these IRM strategies. The first source is current practices used in agriculture but agricultural insecticides tend to have short persistence, target multiple species/stages and must consider the wider ecological impact of their deployment, all of which makes their uncritical adoption into public health problematic (see e.g. Table 1 in [8]). The second source of guidance is empirical field trials but these are expensive, long and operationally challenging (e.g. [9]) and differences in how IRM strategies slow/drive insecticide resistance may be obscured by mosquito movement and gene flow between sites, or the short time frame may be insufficient to allow such differences to become apparent. Their high costs and logistical challenges effectively prevent their replication over transmission settings that may differ widely in vector species, biomics and local details of insecticidal control. The third source of evidence, used here, is to use computer simulation to compare IRM strategies which can be done rapidly, at low-cost, cover a wide range of different deployment settings and scenarios, and can be ‘replicated’ across a wide range of mosquito transmission settings and resistance genetics. This approach was first developed and applied in the 1980s to obtain the basic principles (access to that literature can be found in Tabashnik’s 1989 paper [10]), fell into disuse, and has been recently resurrected to take advantages in recent improvements in computer technology and statistical methods (e.g. [8, 11–14]).This paper is part of that resurrection process.

**Table 1.** Modelling parameters, definitions and distribution ranges used. These are explained in, and mainly taken from *Table 4* of Levick et al [11]. The exceptions are the last two parameters i.e. Simultaneous insecticide contact and Rho. Note that we set “male_exposure_prob” from Levick et al to be 1 which meant males and females have equal probability of contacting insecticides; this was done to simplify the model.

| Variable name | Definition | Parameter distribution |
| --- | --- | --- |
| **Starting frequency allele 1** | Starting frequency of resistant allele at locus 1 and 2 | 0.0001-0.1 (log uniform) |
| **Starting frequency allele 2** |  | 0.0001-0.1 (log uniform) |
| **Exposure** | The proportion of female mosquitoes exposed to the insecticide | 0.1-0.9 |
| **Effectiveness insecticide 1** | Proportion of susceptible (SS) mosquitoes killed after contact with insecticide 1 or 2 | 0.3-1.0 |
| **Effectiveness insecticide 2** |  | 0.3-1.0 |
| **Dominance allele 1** | Dominance of resistance at locus 1 and 2 | 0-1 |
| **Dominance allele 2** |  | 0-1 |
| **rr restoration insecticide 1** | The ability of the resistance (RR) genotype to restore the fitness reduced by exposure to the insecticide | 0.2-1.0 |
| **rr restoration insecticide 2** |  | 0.2-1.0 |
| **Simultaneous insecticide contact (SIC)** | Percentage of insects meeting both panels/insecticides during a feeding cycle. The variation is only relevant for panels because it is set to 1 for mixtures and 0 for micro-mosaics. | 0.2-1.0 |
| **Rho** | Chance of mosquito surviving (i.e., avoiding deaths by natural, non-insecticidal, causes) between blood meals. | 0.3-0.8 |

Three IRM strategies are investigated here on the assumption that two insecticides are ready to de deployed i.e. mixtures, panels and micro-mosaics. The terminology used to describe IRM strategies is highly inconsistent (e.g. [15, 16]), so we briefly describe each strategy to minimise any ambiguity.

Mixtures are formulations that ensure insects will simultaneously contact both insecticides, most likely because they are simultaneously present on all parts of an insecticide-treated bednet (ITN) or are sprayed onto house walls as mixtures in indoor residual spraying (IRS) programmes. Computer simulation studies have consistently identified mixtures as the best-performing IRM strategies, but <u>only</u> if both insecticides are present at concentration sufficiently effective to kill fully susceptible mosquitoes (e.g. [11, 17] but see also [14]). This requirement generates mutual protection i.e. a mosquito resistant to one insecticide would likely be killed by the second insecticide in the mixture, that is the key to mixtures success.

There are two important practical disadvantages to mixtures. First, they are, by definition, more expensive than deploying single insecticides (unless the concentrations of each insecticide are reduced, which runs the risk of lessening the degree of mutual protection). Secondly, physical or chemical properties may preclude their co-application as a mixture. Consequently, alternative strategies for co-deploying two insecticides have been proposed as “pseudo-mixtures”: strategies where mosquitoes are likely, but not guaranteed, to encounter both insecticide during a feeding cycle or lifetime. Foremost amongst these strategies are panel nets (hereafter, “panels”) and micro-mosaics; it is hoped that the two alternatives will be cost-effective ways of obtaining the advantages of mixtures.

Panels. This idea comes from a “panel” ITN where different panels of the same net contain different insecticides (for example the roof panel may contain one insecticide, while the walls may contain another). Operationally, the important factor is that a foraging female mosquito may meet both insecticides in the same feeding cycle; which is equivalent in insecticide exposure terms to her meeting a mixture. In principle it is possible to use the same “Panel” approach in IRS by applying different insecticides to different walls of a house. Alternatively, panels could comprise of one insecticide on the ITN and another insecticide as IRS on the wall of the same hut. [Note that this is often called an insecticide “combination” in the IR literature.]

Micro-mosaics. This strategy deploys different insecticides on a very fine geographical scale, most plausibly by treating adjacent houses in a village with different insecticide. The key difference between panels and micro-mosaics is that mosquitoes <u>may</u> encounter both insecticides during a single feeding cycle if insecticides are deployed as panels in the same ITN or hut, but can only encounter a single insecticide per feeding cycle when deployed as micro-mosaics (we ignore the possibility that a mosquito may forage in one hut then move to another during a feeding cycle). Note that by “micro-mosaic” we mean a mosaic that occurs within a single local breeding population of mosquitoes which is clearly the case here in the “adjacent huts” scenario. We use the prefix “micro-“ to distinguish this strategy from “macro-mosaics” which describes deployments on much larger geographic scales covering different breeding populations, e.g. different insecticides used in different health districts. The potential problem with micro-mosaics is demographic: a female mosquitoes may have produced most of her offspring by the time she meets the second insecticide in a subsequent feed cycle so the impact of the pseudo-mixture effect may be small.

The issue of how best to simultaneous deploy two insecticides has not previously been an issue in public health intervention in resource-poor areas because only a single class, the pyrethroids, was sufficiently affordable and non-toxic for use in these setting. However the success of the Innovative Vector Control Consortium (IVCC) in developing novel or re-purposed active-ingredients and insecticides makes it an important practical question i.e. how best should two insecticides be deployed to minimise selection for resistance? This paper uses computer simulation to compare the effectiveness of mixtures, panels and micro-mosaics and whether the latter two strategies are effective substitutes for a mixture.

## 2. Methods

The main simulation methodology is as described previous by Levick et al [11] and in a more accessible manner by South and Hastings [18].

A key variable in that work was their “correct_mix_deployement” parameter which was used to account for operational issues (such as supply chain failures) that may prevent a mixture being used. The parameter defines how often (in a single feeding cycle) a mosquito meets both insecticides. Those not encountering the “correct deployment” were assumed to meet only a single insecticide with an equal probability of encountering either insecticide. We can therefore use this parameter to investigate the three strategies. Setting correct_mix_deployment to 1.0 investigates mixtures because mosquitoes, by definition, must encounter both insecticides. It takes a value of zero for micro-mosaics because, by definition, mosquitoes never meet both insecticides during a feeding cycle (we ignore the possibility of interrupted feeding when a mosquito may encounter one insecticide in a hut, fly to an adjacent one and met the other insecticide). For panels, the parameter defines how often a mosquito meets both panels/insecticides and was allowed to vary between 0.2 and 1.0 with the assumption that if the mosquito does encounter only one insecticide is has equal probabilities of meeting each insecticide. The name “correct_mix_deployment” is slightly opaque in the current context so we rename it in this manuscript to “Simultaneous insecticide contact” (SIC).

There is an implicit assumption in the Levick et al model that a mosquito has only one opportunity to encounter insecticide(s) before breeding, equivalent to there being only one foraging/oviposition cycle per generation. This is, as far as we are aware, the same in all population genetic models of insecticide resistance (the only exception we know is by Read and colleagues [19], based on the transmission model of [20]). This assumption has to be removed to investigate micro-mosaics where a key argument for their deployment is that mosquitoes may meet different insecticides in different foraging cycles, allowing micro-mosaics to have some/most of the benefits of mixtures, but at reduced deployment costs.

It is relatively straightforward to incorporate multiple gonadotrophic cycles into the methodology of Levick et al [11]. That work relied on defining the fitnesses of 18 genotypes (9 resistance genotypes by 2 sexes) such as W^m,SS1SS2^ which is the fitness of males with resistance genotype SS1SS2 (i.e. homozygous sensitive at both loci), W^f,SS1SR2^ which is the fitness of females with resistance genotype SS1SR2 (i.e. homozygous sensitive at locus 1 but heterozygous at locus 2), and so on. These fitnesses were the probability of surviving possible contact with the insecticides in a <u>single</u> feeding cycle. So the probability of surviving and reproducing in the first feeding cycle is W*ρ
Where

- W is fitness i.e. the probability of surviving possible insecticide contact (e.g. _W_f,SS1SR2 _)_
- ρ is probability of surviving death by natural causes (i.e. not associated with insecticide contact) during the feeding cycle.

We did not require ‘ρ’ in the Levick et al approach because multiplying all values of W by ‘ρ’ would not alter the relatively sizes of the 18 fitness values.

The probability of surviving and reproducing during the second feeding cycle is [Wρ]*Wρ

Where the term in square brackets is the probability of surviving the first cycle which is obviously a prerequisite for reproducing in the second.

The probability of surviving and reproducing during the third feeding cycle is [Wρ*Wρ]*Wρ

Where the term in square brackets is the probability of surviving the first two cycles. And so on for all feeding cycles. The total reproductive output over all feeding cycles, denoted *W^^^*, is simply the sum of each feeding cycle i.e.

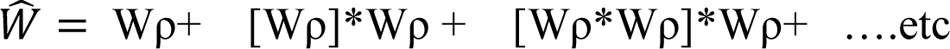

Which is

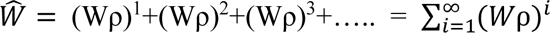

There is algebraic result that allows this sum to be easily calculated i.e.

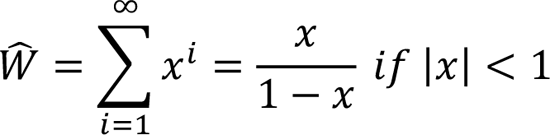

Where x= W*ρ for algebraic clarity

So to investigate multiple feeding cycles we first calculate the 18 fitness values of W as already described by Levick et al and then simply update them to

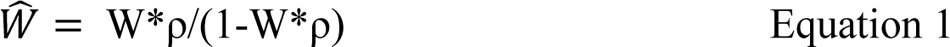

So Equation 1 of Levick et al [11] would be updated as follows

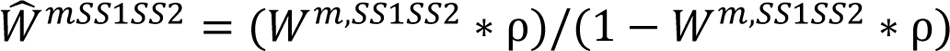

Computationally, we simply update the fitness of the 18 genotypes from W to *W^^^* as described above. This requires editing 18 lines of code and then using the existing code described in Levick et al [11] and South and Hastings [18].

There are two implicit assumptions built into this approach. Firstly that females mate once, before their first egg lay, and store sperm for subsequent egg laying. This appears to be the case for Anopheline mosquitos and means we do not have to consider different mating combinations in each gonadotrophic cycle. Secondly, we have made the argument in terms of females egg laying cycles and this obviously does not apply to males who enter mating swarms rather than laying eggs. In the case of male their “reproductive output” is their number of successful matings rather than number of eggs laid but exactly the same argument applies. It is unclear whether males enter a mating and swarm remain until death, or whether they enter a swarm then leave to recover before re-joining. The calculations above assume the latter and that males’ “swarming” intervals are the same as the females egg laying intervals. It would be simple to specify a different interval for males, but we kept male and female intervals identical here for simplicity and to avoid adding additional parameters to the model.

The model assumes only two novel insecticides are available, and therefore, a failing insecticide cannot be replaced with a third insecticides to combat resistance. The starting assumption is that there is resistance to each insecticide in the population but at very low frequencies. All insecticides are fully active for the product’s lifetime, and the product is replaced at the end of its deployment period (usually every year for IRS and every three years for ITNs). Again, this is a common assumption in modelling IR but there is increasing interest in how insecticide effectiveness declines after deployment (see [12] for discussion and access to this literature).

We assume independent mechanisms of resistance for each insecticide i.e. different genes encode resistance to each insecticides and there is no cross-resistance. The effects of insecticides in a mixture are multiplicative, e.g. if the probability of surviving exposure to insecticide A alone is 0.3 and of surviving insecticide B alone is 0.2, then the probability of surviving expose to a mixture of A and B is 0.3*0.2 = 0.06. No synergistic or antagonistic interactions between insecticides will be explored in this time frame.

These model assumptions are basically those of most population-genetic models i.e. that individual mosquitoes mate at random, have non-overlapping generations and Mendelian genetic inheritance. The model assumes deployment scenarios are all within the same mosquito breeding population with no emigration or immigration.

The key output for evaluating the alterative IRM strategies is “Time to resistance” which is defined (as previously e.g. [11, 17]) as the time, in generations, at which resistance allele frequency (RAF) exceeds 50% for both insecticides.

The simulations were run for 500 generations and analysed at 250 and 500 generations. Most anopheline mosquitoes (the vectors of malaria) have around 10 generations per year in highly endemic regions so 250 generations equate to around 25 years which is a common time horizon for operational and economic planning. However, some simulations had not reached the endpoint (RAF>50%) by 250 generations so running for the full 500 generations allow simulations more opportunity to detect a difference between the strategies and are better for sensitivity analysis. We compare the performances of each strategy as the ratio of the times until resistance is reached (see caption to Figure 1). If one strategy (but not the other) has still not reached resistance by end of the simulation (250 or 500 generations) we use that figure, i.e. 250 or 500, as time for resistance when calculating the ratio; this allows us to use information from that run, but means we under-estimate the performance of the better strategy. Each simulation run is based on a random sample of the parameter space given in Table 1 and each strategy (mixture, panels, micro-mosaic) was run under the same parameter set and times to resistance measured for each strategy. We ran 5000 simulations to compare strategies under the parameter space given in Table 1.

**Figure 1.**
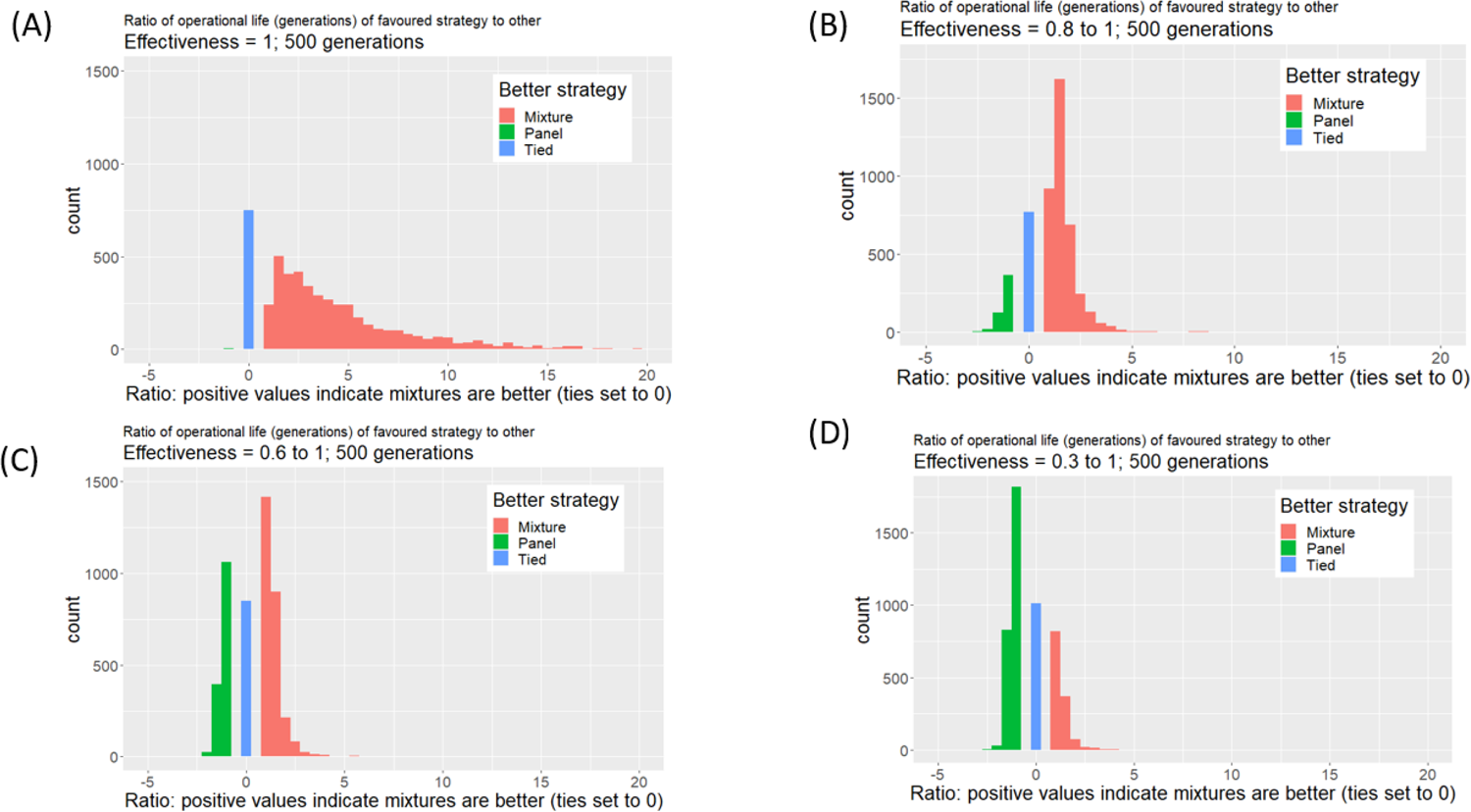
A comparison of Mixtures vs Panels as IRM strategies when each strategy was tested under the same parameter set (see Table 1) and the simulation run for 500 generations, or until resistance has arisen in each strategy (i.e. RAF >0.5 for both insecticides). The ratio of time till resistance arises was calculated as time(better strategy)/time(worse strategy) and we set the ratio to be positive when Mixtures perform best, and negative when Panels perform best. This was done for 5000 parameter combinations and results shown on the histogram below. Runs where mixture perform best are shown in red and runs where panels performed best are shown in green. The Xaxis indicate the magnitude of the advantage; for example, a value of +10 means mixtures last 10 times longer than panels while a value of −5 means panels last 5 times longer than mixtures. Panel (A): Effectiveness =1, Panel (B) Effectiveness varies from 0.8 to 1.0 Panel (C) Effectiveness varies from 0.6 to 1.0 Panel (D) Effectiveness varies from 0.3 to 1.0

The simulations were all run in R Statistical Software (v4.2.2 [21]) through Rstudio v2022.12.0 [22]. The code for running the simulation is as described in Levick et al [11], and is available as “runModel2” from a Github repository at https://github.com/andysouth/resistance/blob/master/R/runModel2.r. PRCC analysis was performed using the function “pcc” (with rank=TRUE) from the library “ppcor” [23]. Classification trees were produced and plotted using the functions “rpart” and “rpart.plot” (from the packages rpart and rpart.plot) and included all the variables listed on Table 1.

## 3. Results

Figure 1 shows the results from a comparison of mixtures vs panels. Effectiveness of the insecticides (i.e. ability to kill SS mosquitoes) is a key factor so we present 4 comparisons i.e. when effectiveness is set to 1.0 and when effectiveness is allowed to vary between 0.8 to 1, 0.6 to 1 and 0.3 to 1.0. As effectiveness is allowed to fall the advantages of mixtures declines and once effectiveness is allowed to vary between 0.3. and 1, the use of mixtures may become counterproductive, and a Panel strategy is, on average, better (Figure 1D). The PRCC analyses of these results is given on Figure 2. If effectiveness is set to 1 (Figure 1A), then the most important factor is exposure: increasing exposure driving increased advantage of mixtures over a panel strategy. A second factor, simultaneous insecticide contact (SIC) has a smaller impact with increased SIC being associated with reduced advantage of mixtures over panels: this is as expected because as SIC increases then mosquitoes are more likely to encounter both types of panels/insecticides so panels behave more like a mixture. Setting effectiveness to one is an idealised case and Figure 2B presents PRCC when effectiveness is allowed to vary between 0.3 and 1. As expected from Figure 1, effectiveness is the largest factor determining whether mixtures are the best policy with increasing effectiveness driving an increased probability that mixtures outperform panels. Exposure also plays a role when effectiveness declines, decreasing exposure favouring mixtures. Interestingly, as in Figure 2A, correct mix deployment also has an impact but in the different direction i.e. an increase in its value favours mixtures as a strategy.

**Figure 2.**
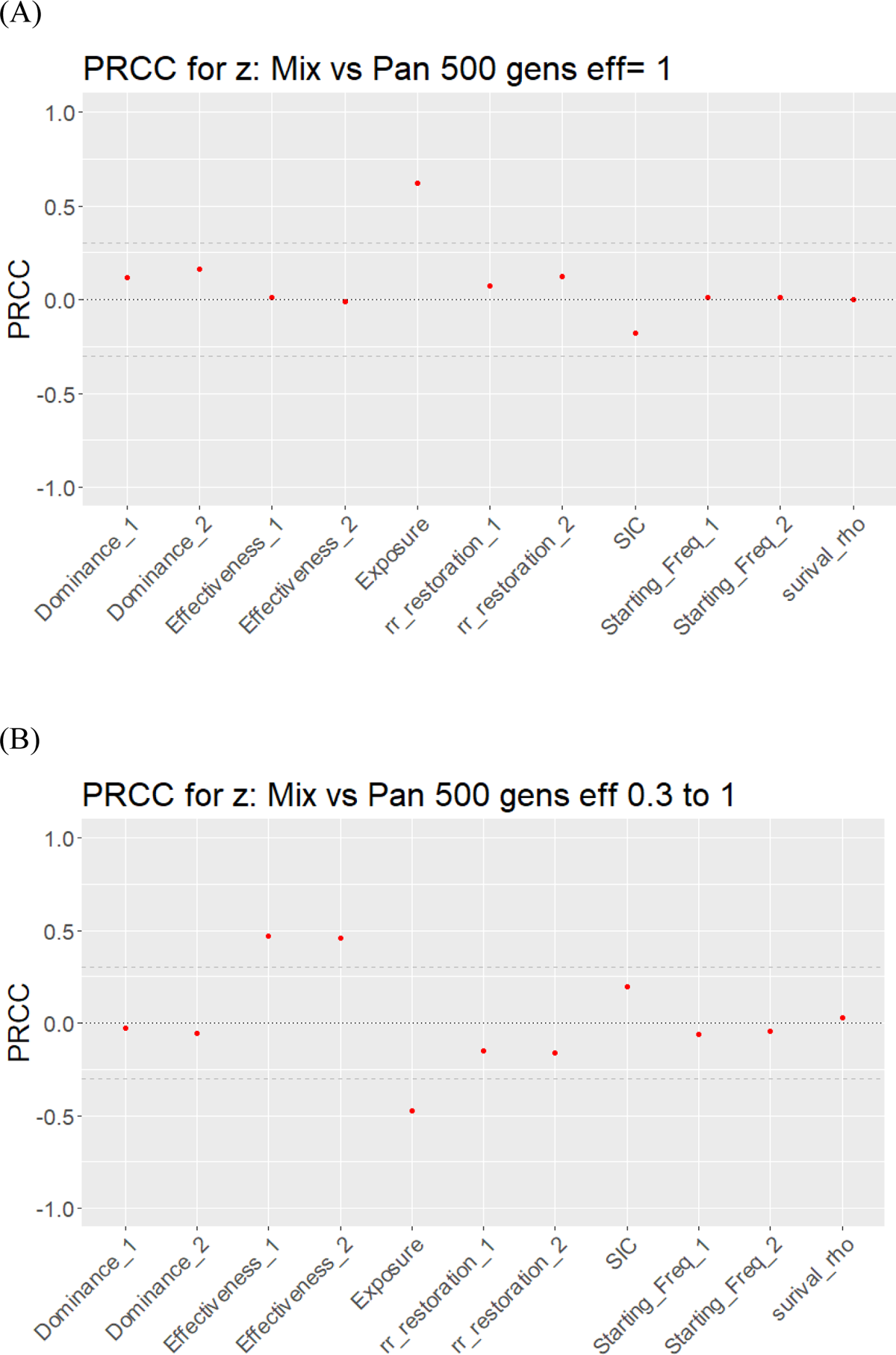
Partial Rank Correlation Coefficients (PRCC) for comparison of Mixtures vs Panels as IRM strategies as measured by the ratios of their time to resistance. Panel (A) assuming Effectiveness is 1 (i.e. analysing the data shown Figure 1A). Panel B assuming Effectiveness varies between 0.3 and 1 (i.e. analysing the data shown in Figure 1D).

Figures 3 and 4 are analogous to Figures 1 and 2, but present and analyse the comparison of mixtures against micro-mosaic. Interestingly, the results are almost identical (noting that SIC has no impact in Figure 4 (unlike Figure 2) because the parameter only applies to Panels which are not included in the comparison). Notably, the survival parameter, rho, has no impact on whether mixtures or micro-mosaics were better. This is slightly surprising because the putative advantage of micro-mosaics is that mosquitoes may meet different insecticides in subsequent feed cycles and increased survival probability during feeding will increase this chance.

**Figure 3.**
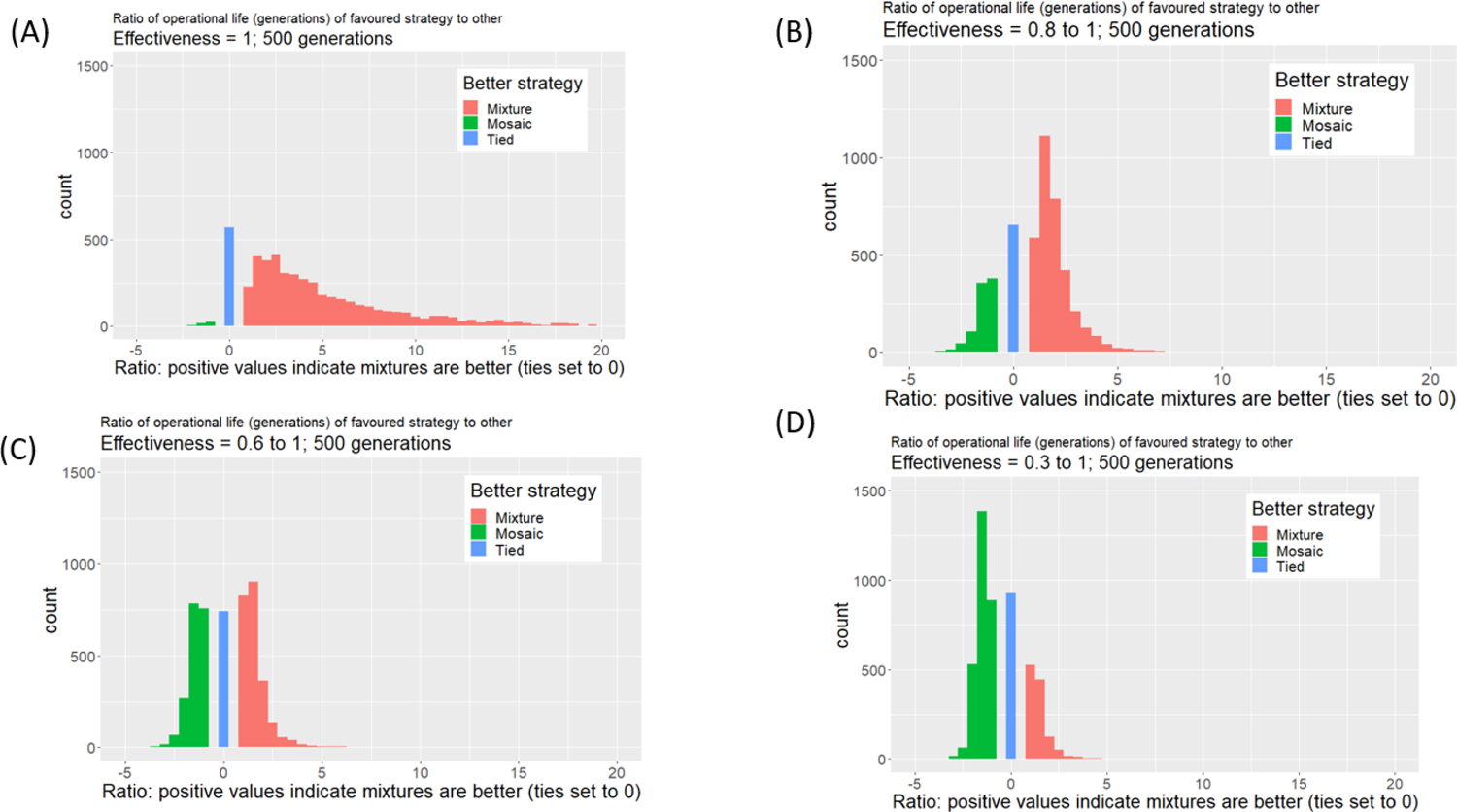
As for Figure 1 except the comparison is between Mixtures and Micro-mosaics as IRM strategies.

**Figure 4.**
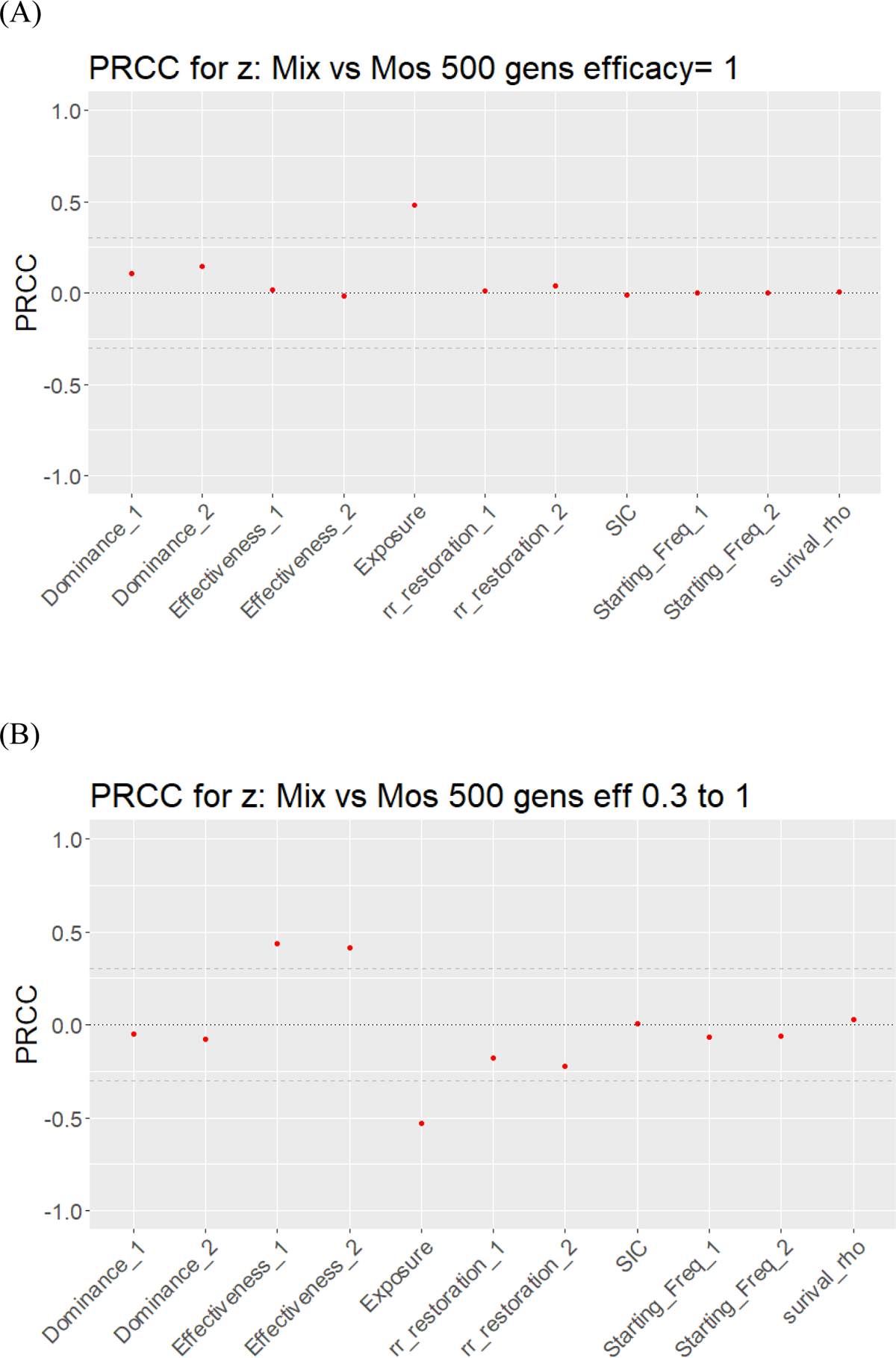
As for Figure 2 except that this figure shows PRCC analysis of mixtures against mosaics. Panel (A) assuming efficacy is 1 (i.e., analysing the data shown in Figure 3A). Panel B assuming efficacy varies between 0.3 and 1 (i.e., analysing the data shown in Figure 3D). Note that SIC has no impact in this analysis as it is always set to 1 for mixtures and 0 for micro mosaics (it is included to keep PRCC plot structures consistent).

Figures 5 and 6 are analogous to Figures 1 to 4 but show results from a comparison of Panel vs micro-mosaics. Recall that panels are intermediate between mixtures and micro-mosaics (they only differ in the SIC parameter which is 0 in mosaics, 1 in mixtures and varies in panels; see table 1). Effectiveness therefore would be expected to have a large effect in favouring panels: this occurs but is much smaller than in mixtures (i.e. Figure 5 vs Figure 1) with Panels being slightly more favourable when effectiveness is 1 (Figure 5A) but declining to the extent than micro-mosaics are favoured as effectiveness falls. Previous analyses shows that when effectiveness is 1 then SIC is a key factor, presumably because as its value increases the panels behave more like mixtures. Coverage is also a factor i.e. reducing coverage favours Panels. When Effectiveness is allowed to vary between 0.3 and 1 the PRCC analysis is virtually identical to that obtained from the other two comparisons (i.e. Figure 6B vs Figure 2B and 4B).

**Figure 5.**
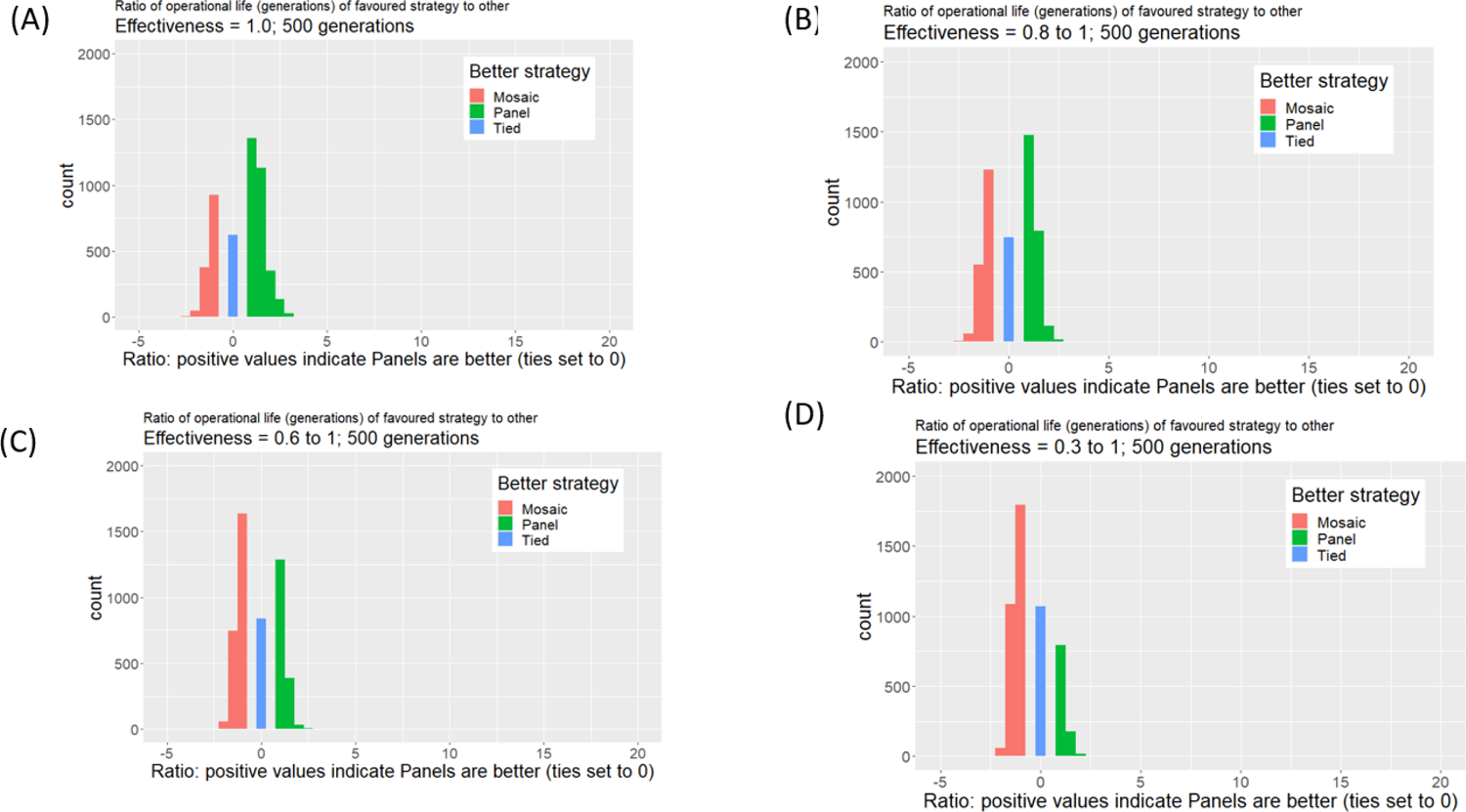
As for Figures 1 and 3 except the comparison is between Panels and Micro-mosaics as IRM strategies.

**Figure 6.**
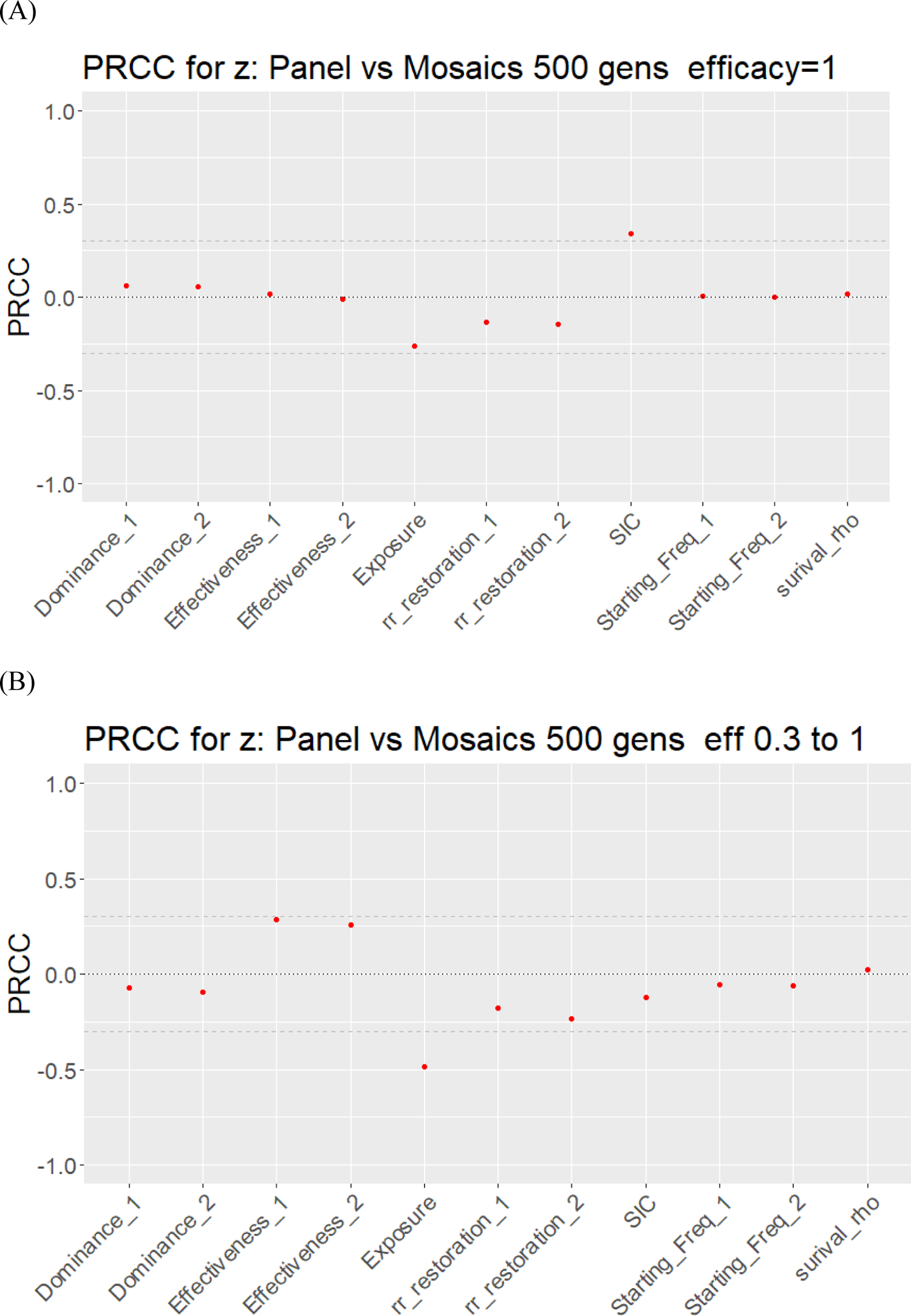
As for Figures 2 and 4 except that this figure shows PRCC analysis of Panels against micro-mosaics. Panel (A) assuming efficacy is 1 (i.e. analysing the data shown in Figure 5A). Panel B assuming efficacy varies between 0.3 and 1 (i.e. analysing the data shown in Figure 5D).

The PRCC analysis are informative but report the impact of each variable independently. Classification trees may increase understanding through examining variables in combination, with the hope that some parameter combination may clearly favour one strategy over another. One decision to make in constructing the trees is the value of the complexity parameter (cp): a high value allows detailed trees to be constructed but with the penalty that they may become so detailed as to be useless in guiding decisions. A higher value of cp results in less precision (i.e. more misclassifications) but the trees are simpler and may identify parameter space of sufficient breadth that they could be used for policy decisions. We therefore set cp=0.02 and the resulting Classification trees are shown on Figure 7 to 9. They are consistent with the results from PRCC i.e. that effectiveness is the key variable but, in our opinion, do not identify combinations of parameters that are sufficiently robust to clearly recommend one strategy over another.

**Figure 7.**
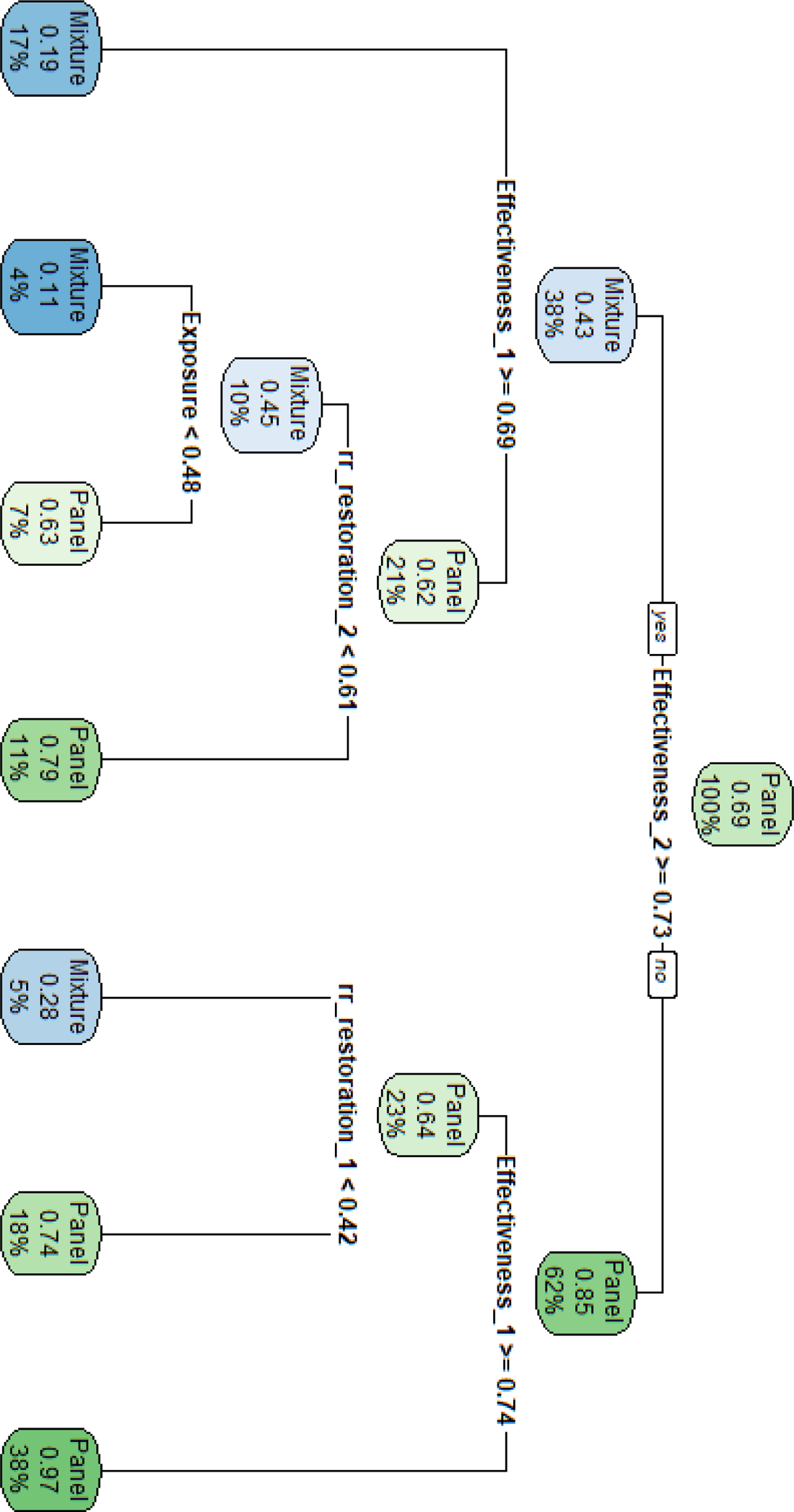
Classification tree for a comparison between Mixtures vs Panels.

**Figure 8.**
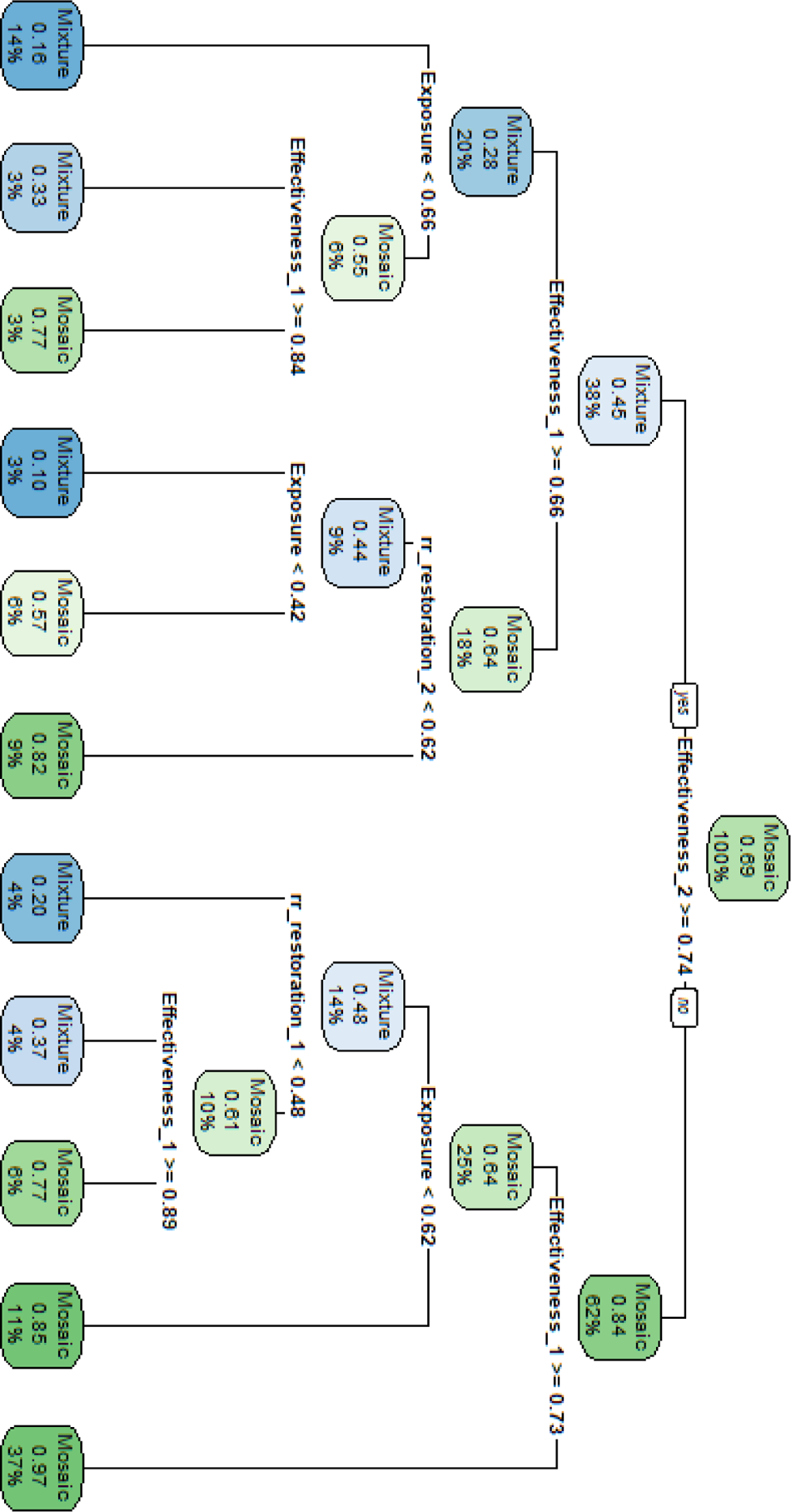
Classification tree for a comparison between Mixtures vs Micro-mosaics.

**Figure 9.**
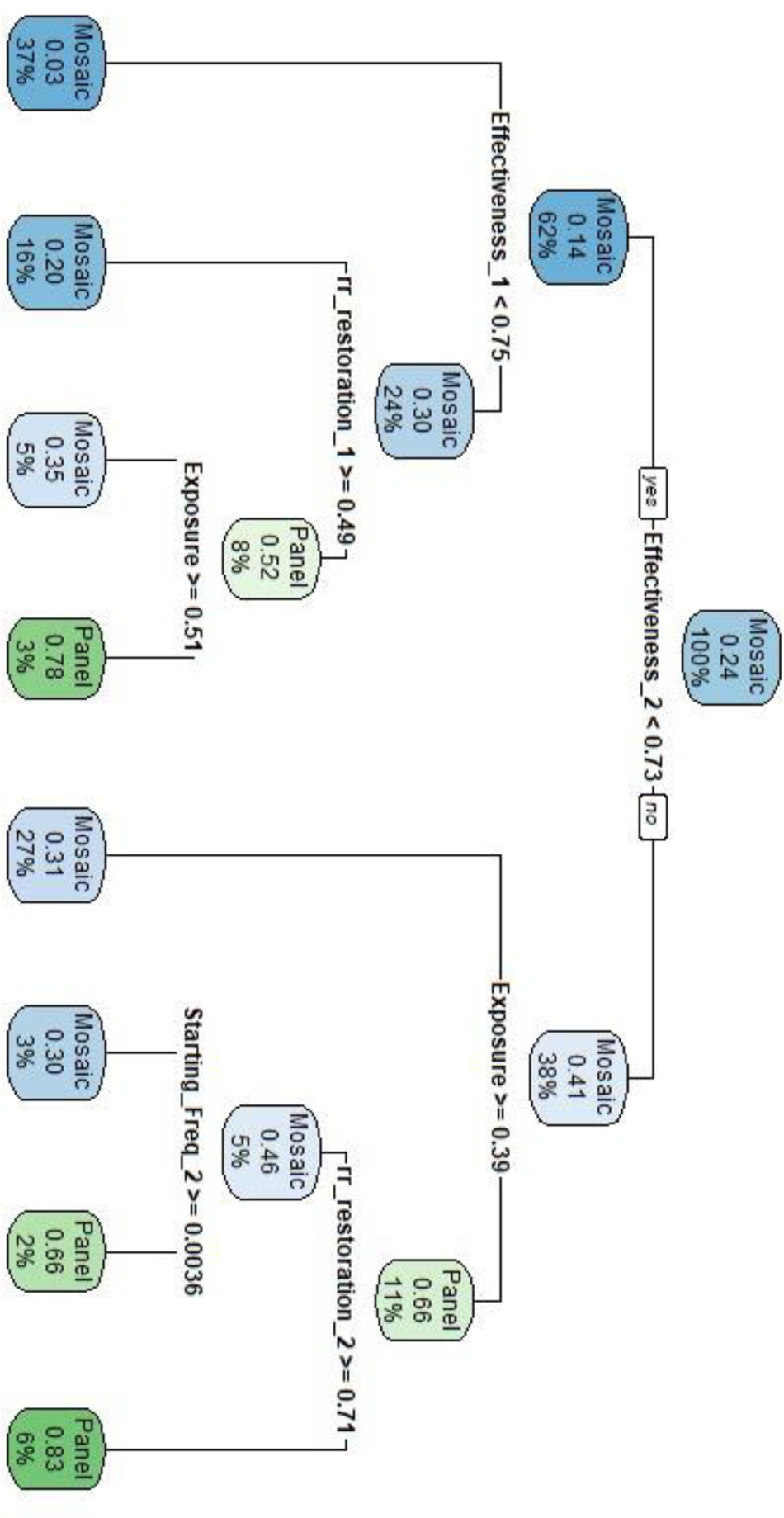
Classification tree for a comparison between Panels. Vs micro-mosaics

The results in Figures 1 to 9 were obtained from simulations run for 500 mosquito generations (roughly 50 years) with this timeline chose to improve detection of differences between strategies (some may still have been tied at earlier generations). A 250 generation (i.e. 25 year) timeline may be more relevant for operational planning. Plots analogous to Figures 1 to 9, but obtained using results at 250 generations, are given in Supplementary Information #1: as can be seen, the results were virtually identical.

## 4. Discussion

The most disappointing results was the poor performance of micro-mosaics as a IRM strategy. Advocates of micro-mosaic nets have argued that foraging females may meet different insecticides in different feeding cycles, allowing micro-mosaics to achieve many of the benefits of mixtures at a reduced cost. However, mortality among foraging females is likely to be high in natural settings so few female mosquitoes actually survive sufficiently long to meet both insecticides, and our results suggest micro-mosaics are not an effective IRM strategy and that they routinely, and substantially, underperform compared to fully-effective mixtures (Figure 1A). Importantly, they start to perform equivalently, or even out-perform, mixtures if the insecticides are not fully effective at killing sensitive mosquitoes (Figures 1 and 2). It is already known that mixture with reduced effectiveness of constituent insecticides offer little, if any, advantage over sequential deployment (e.g. [11, 18] although Madgwick and Kanitz [14] present analyses more favourable to mixtures) and it is likely that the same conclusion will apply to micro-mosaics compared to sequential deployment (we did not simulate sequential deployment as a baseline comparator).

A similar argument applies to Panels which, as for micro-mosaics, have been proposed as a low-cost alternative to Mixtures. Panels perform worse than mixtures when effectiveness is very high, but panels may be better than Mixtures when effectiveness is less than 1.

When Panels and compared to micro-mosaics (Figure 5) there was little difference between the two strategies.

The results highlight, unsurprisingly in view of previous work (e.g. [11, 18, 24]), that mixtures need to maintain the effectiveness of their constituent insecticides over their lifetime both to reduce the selection for resistance, and to maintain their advantage over panels and micro-mosaics as an IRM. The main threats to continued effectiveness are physical damage (when deployed on ITNs) and natural degradation of insecticide activity [12] which is likely to apply to both ITNs and IRS.

The results can therefore be easily understood with reference to previous analyses which showed that reduced effectiveness of insecticides (i.e. reduced ability to kill SS genotypes) in a mixture starts to drastically reduce the advantages of mixtures as a IRM. The reason is that fully effective insecticides provide strong mutual protection i.e. a mosquito resistant to one insecticide will almost certainly be killed by the second insecticide (provided it is sensitive to that insecticide). Panels potentially allows mosquitoes to contact both insecticides, but our parameter SIC recognises than some mosquitoes may not meet both panel types: this reduces the mutual protection of insecticides potentially offered by panels which makes them inferior to fully effective mixtures. An analogous explanation applies to micro-mosaics: by the time a second insecticide can potentially kill a mosquito resistant to the first insecticide the mosquito will likely have laid a large proportion of her eggs (Figure 10) substantially reducing the potential advantages of mutual protection of co-deployment in a micro-mosaic. If effectiveness and coverage are both high then fitness of sensitive mosquitoes will be low and virtually all their eggs will be laid in the first or second oviposition cycle. The relative merits of micro-mosaics vs panels, would therefore depend on their relative loss of mutual protection either spatially (the SIC parameter in panels) or temporally (the mosquito survival rate in micro-mosaics).

**Figure 10.**
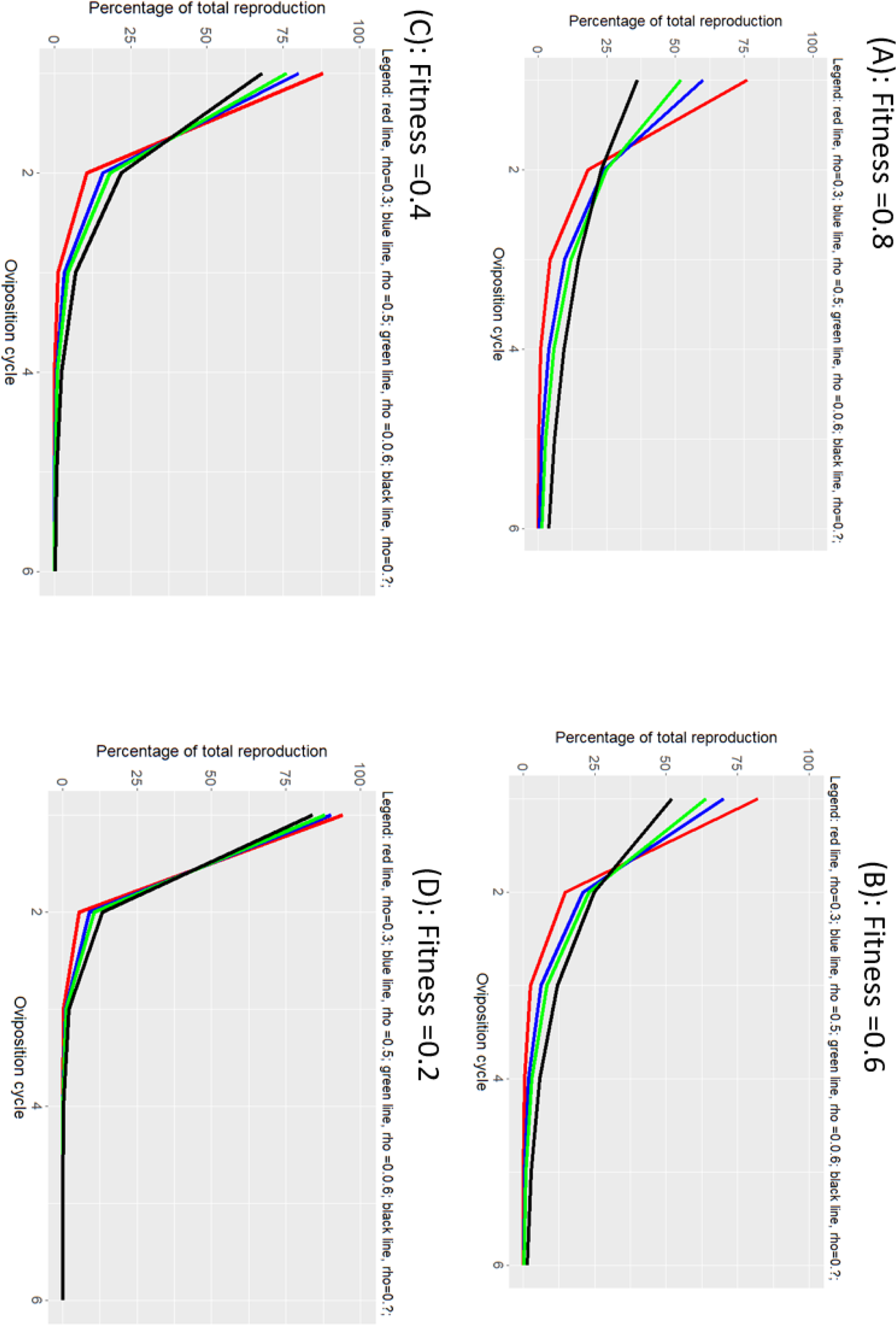
The relative reproduction over successive oviposits from female mosquitoes. This is important for micro-mosaics where part of their attraction is the possibility that mosquitoes may meet different insecticides in different oviposition cycles, thereby gaining some of the benefits of a mixture. The magnitude of oviposits each cycle depend on survival between oviposits as described in the derivation of Equation 1. The fitness of a mosquito genotype depends on its ability to survive insecticide contact and the probability that it meets insecticide (i.e. insecticide coverage). Fitness therefore differs depending on genotype and coverage so we present result for four different fitness with rho (the “natural” survival over a feeding cycle in the absence of insecticide) varied between 0.2 and 0.8 (Table 1).

In summary, if insecticide effectiveness is high (a known prerequisite for mixtures being an effective IRM strategy [18]), then neither Panels nor Micro-mosaics appear to be an effective alternative to mixtures, the latter typically having a 2 to 10 fold increase in time to resistance compared to these alternatives. Once effectiveness falls below 1 then it is difficult to predict which of the three strategies will perform better so no clear recommendations can be made on the basis of their IRM properties and deployment decision will most likely be based on other factors such as cost, availability and operational simplicity.

## Supporting information

SI #1. Analysis restricted to 250 generations

