## Supplementary material for "How best to co-deploy insecticides to minimise selection for resistance": SI #1. Analysis restricted to 250 generations

### Supplementary Information #1: Re-analysis of data restricted to 250 generations.

The main text presents analysis of resistance spread during the 500 mosquito generations after deployment of insecticides Most mosquito vectors of malaria have around 10 generations per year, so this equates to a 50 year time horizon.

Most economic and operational decisions are based on 25 year timelines so this Supplementary Information shows results when analysis is restricted to the first 25 years (250 mosquito generations) post-deployment.

Figure S1. As for Figure 1 of the main text, but analysis based on 250 mosquito generations post-deployment.

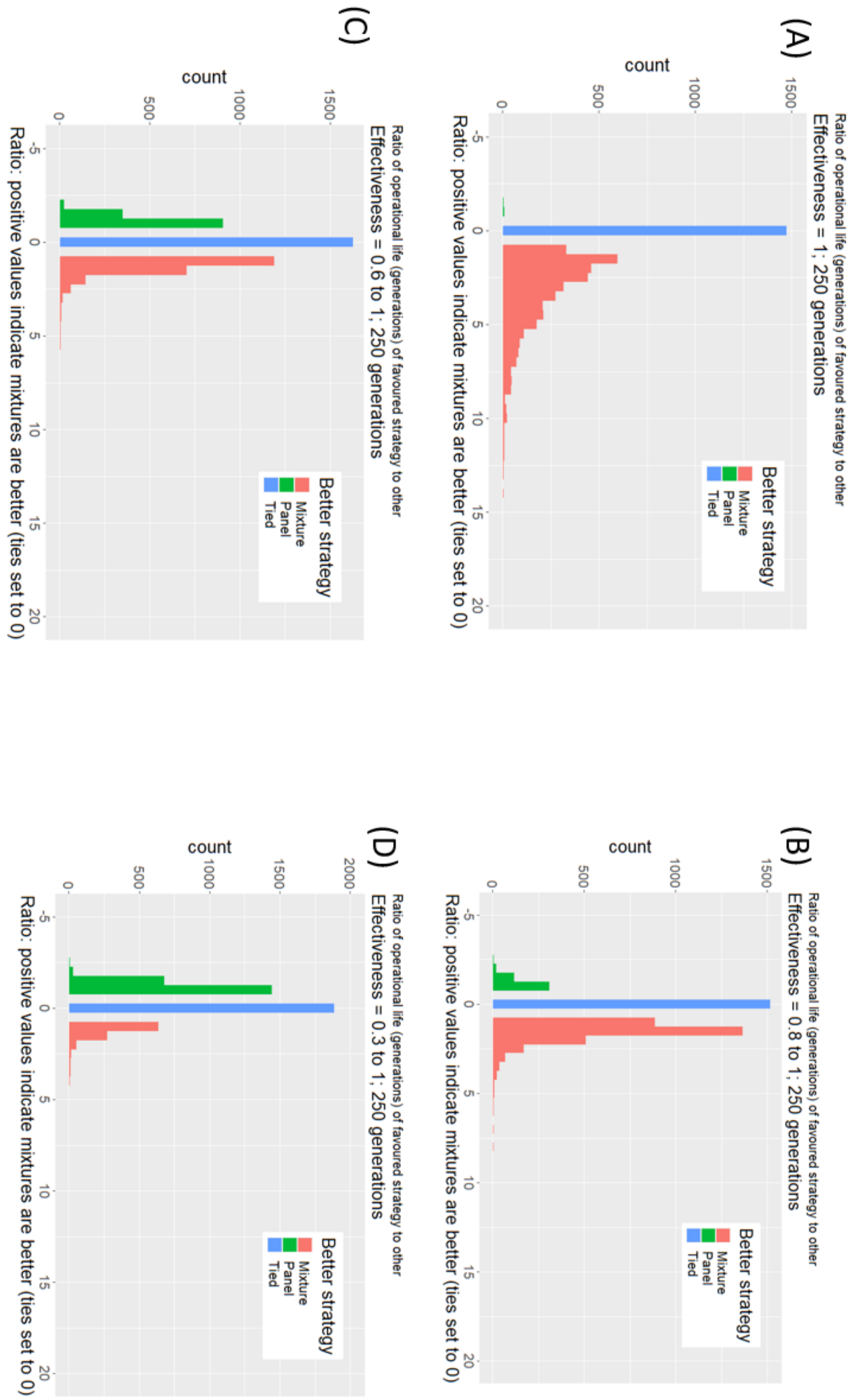

Figure S2. As for Figure 2 of the main text, but analysis based on 250 mosquito generations post-deployment.

(A)

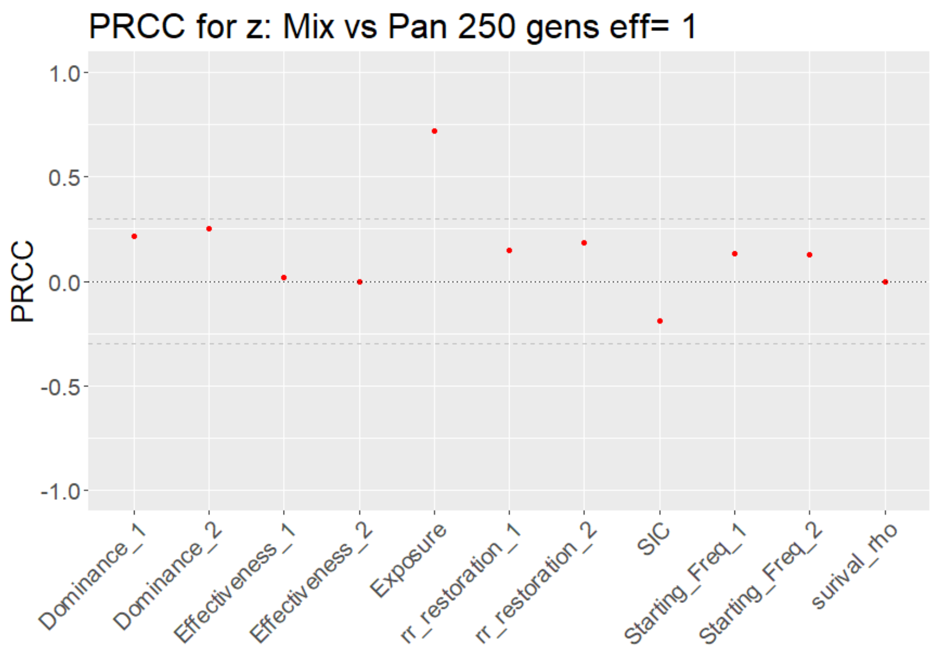


(B)

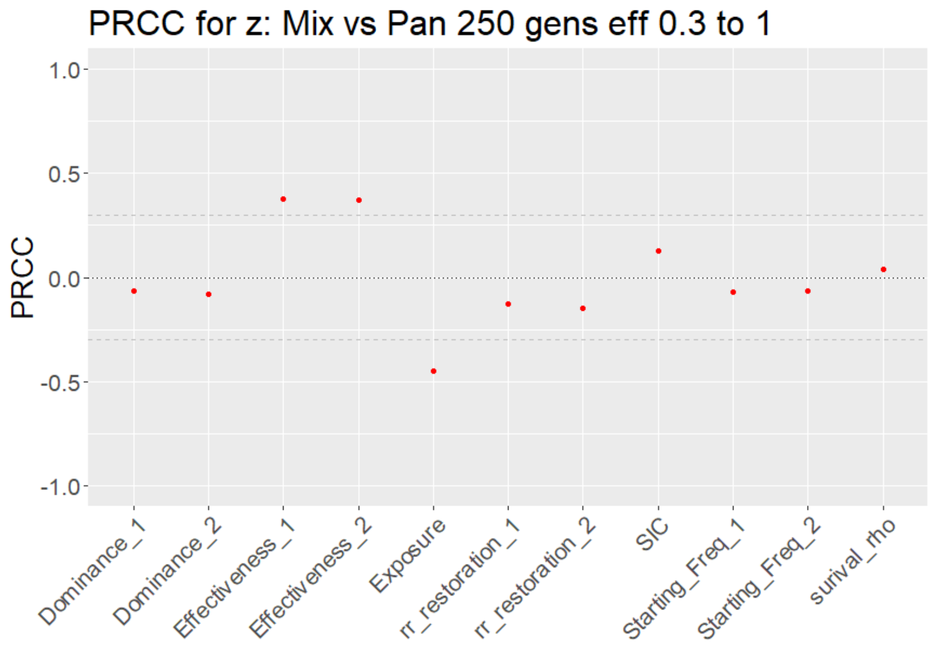

Figure S3.  As for Figure 3 of the main text, but analysis based on 250 mosquito generations post-deployment.

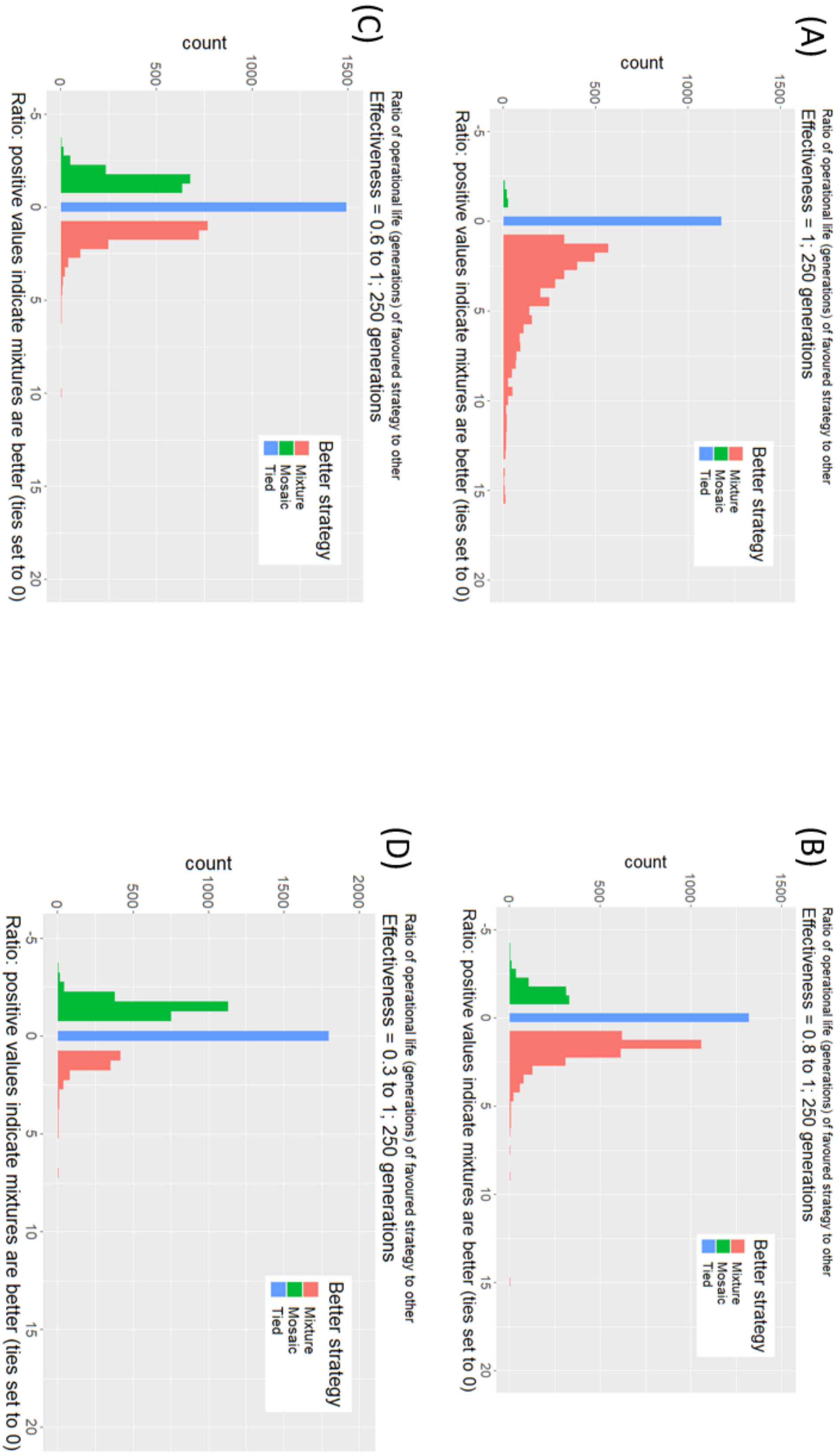

Figure S4. As for Figure 4 of the main text, but analysis based on 250 mosquito generations post-deployment.

(A)

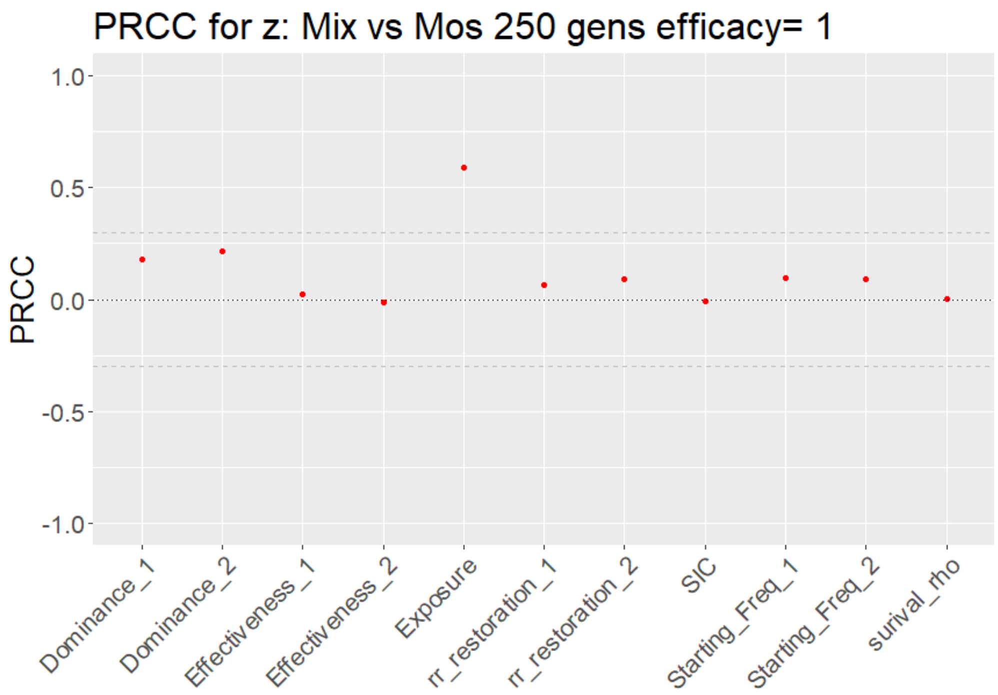


(B)

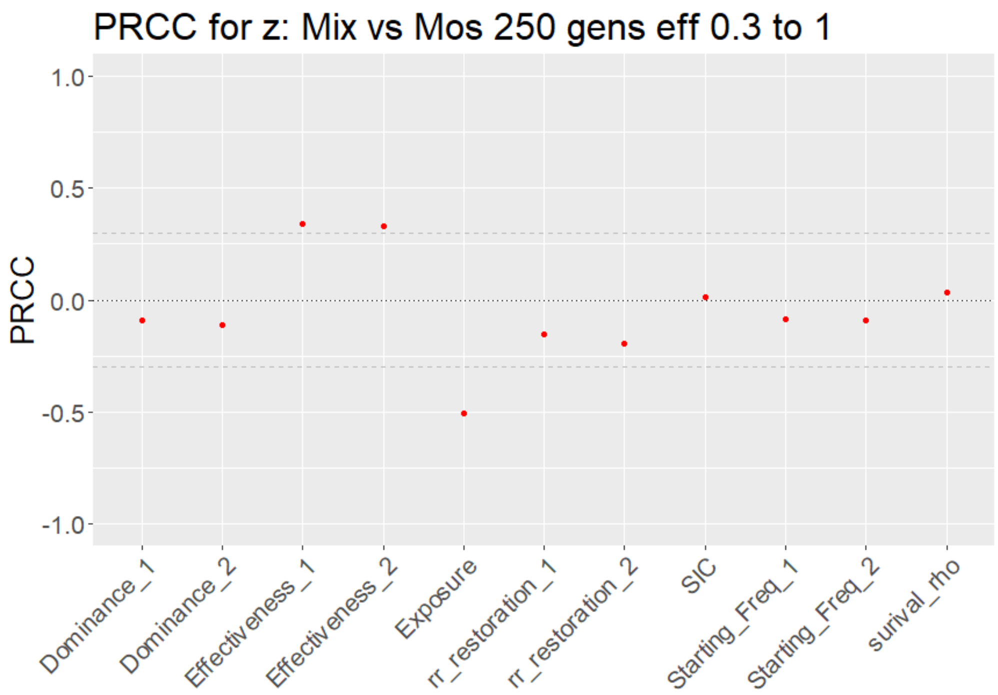

Figure S5. As for Figure 5 of the main text, but analysis based on 250 mosquito generations post-deployment.

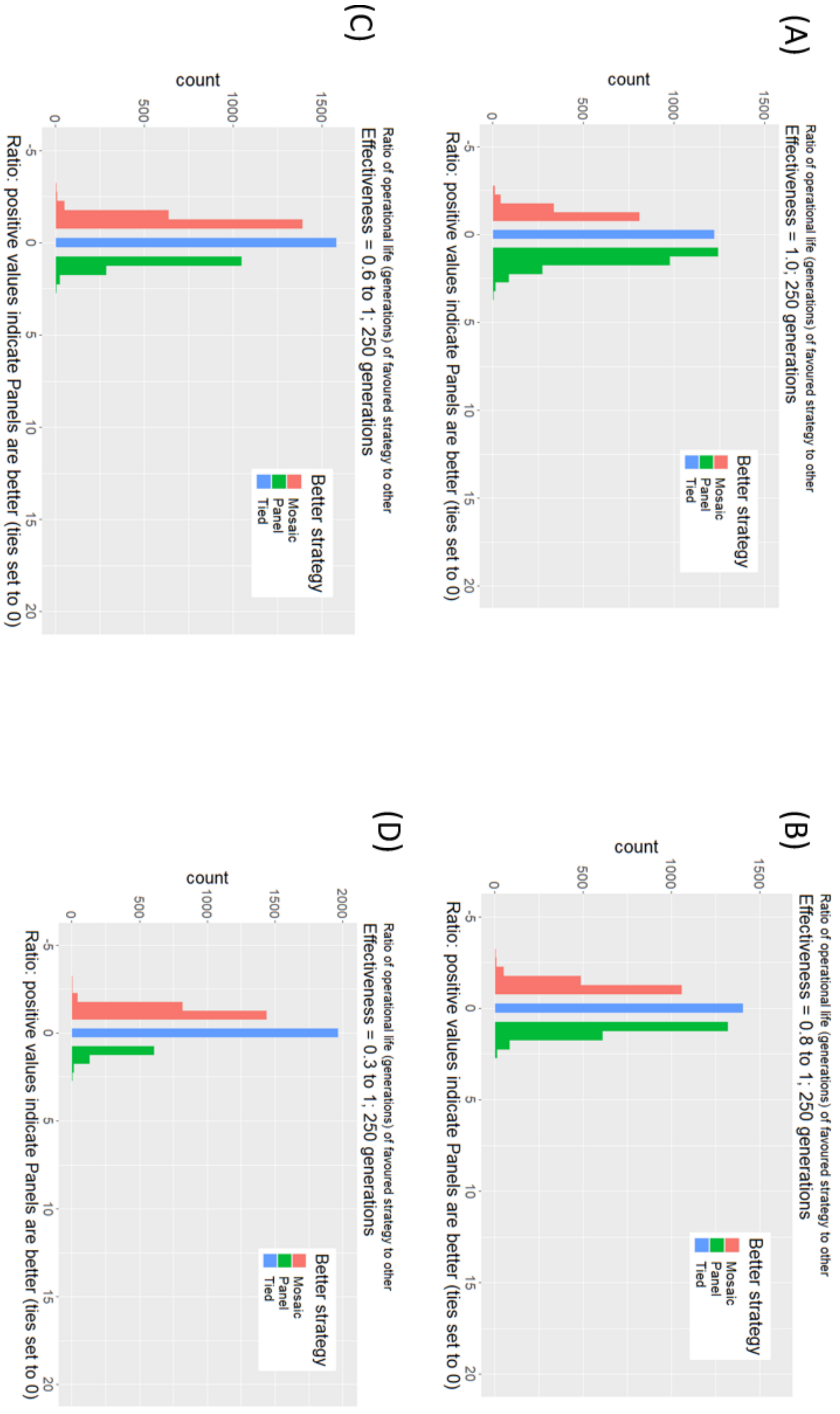

Figure S6. As for Figure 6 of the main text, but analysis based on 250 mosquito generations post-deployment.

(A)

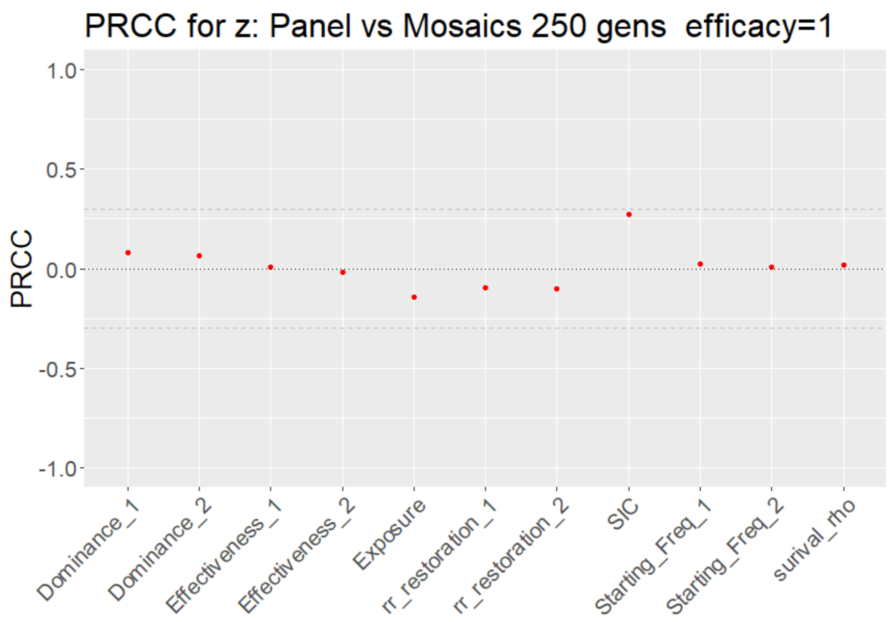


(B)

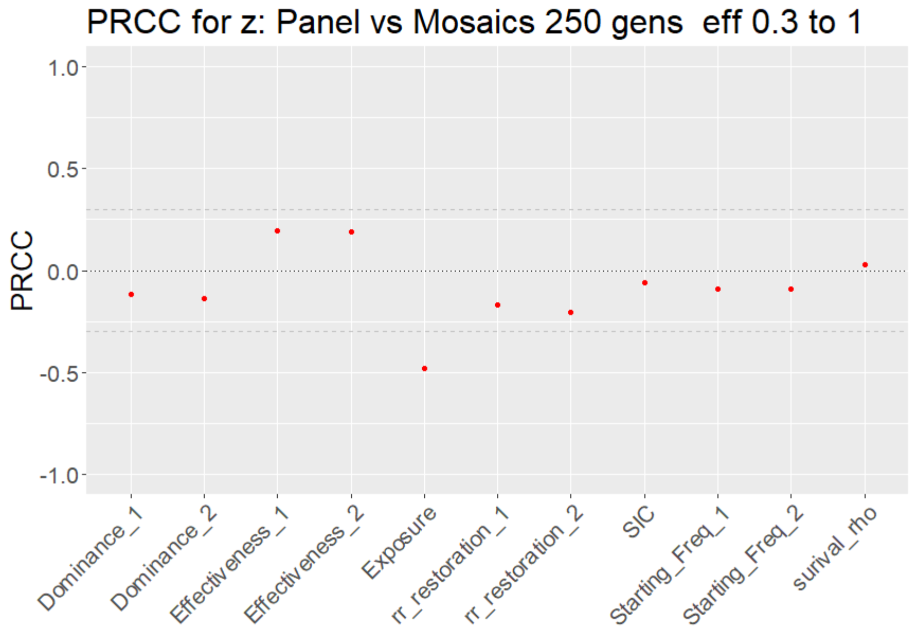

Figure S7. As for Figure 7 of the main text, but analysis based on 250 mosquito generations post-deployment. Classification tree is for a comparison between Mixtures vs Panels.

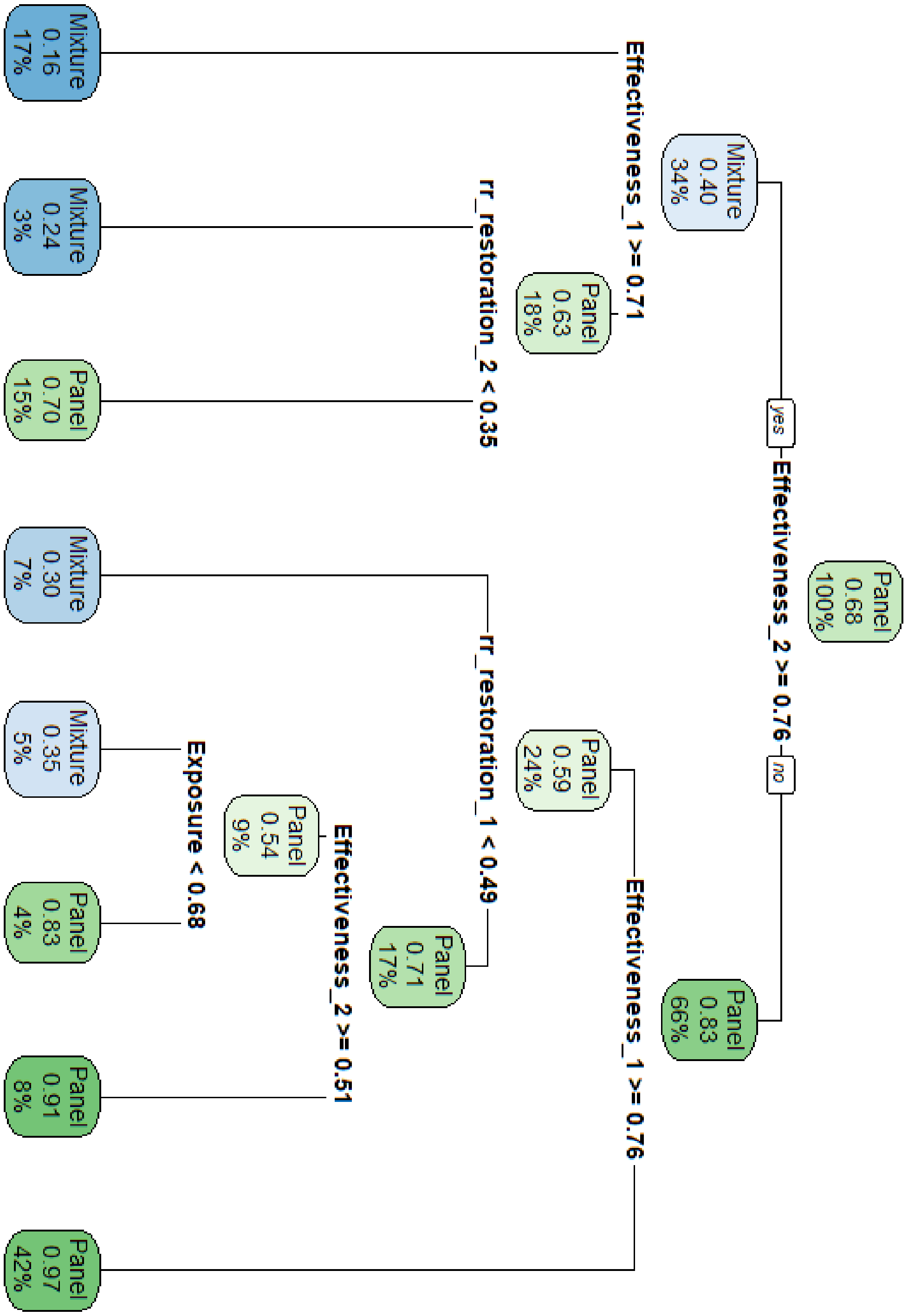

Figure S8. As for Figure 8 of the main text, but analysis based on 250 mosquito generations post-deployment. Classification tree is for a comparison between Mixtures vs Micro-mosaics.

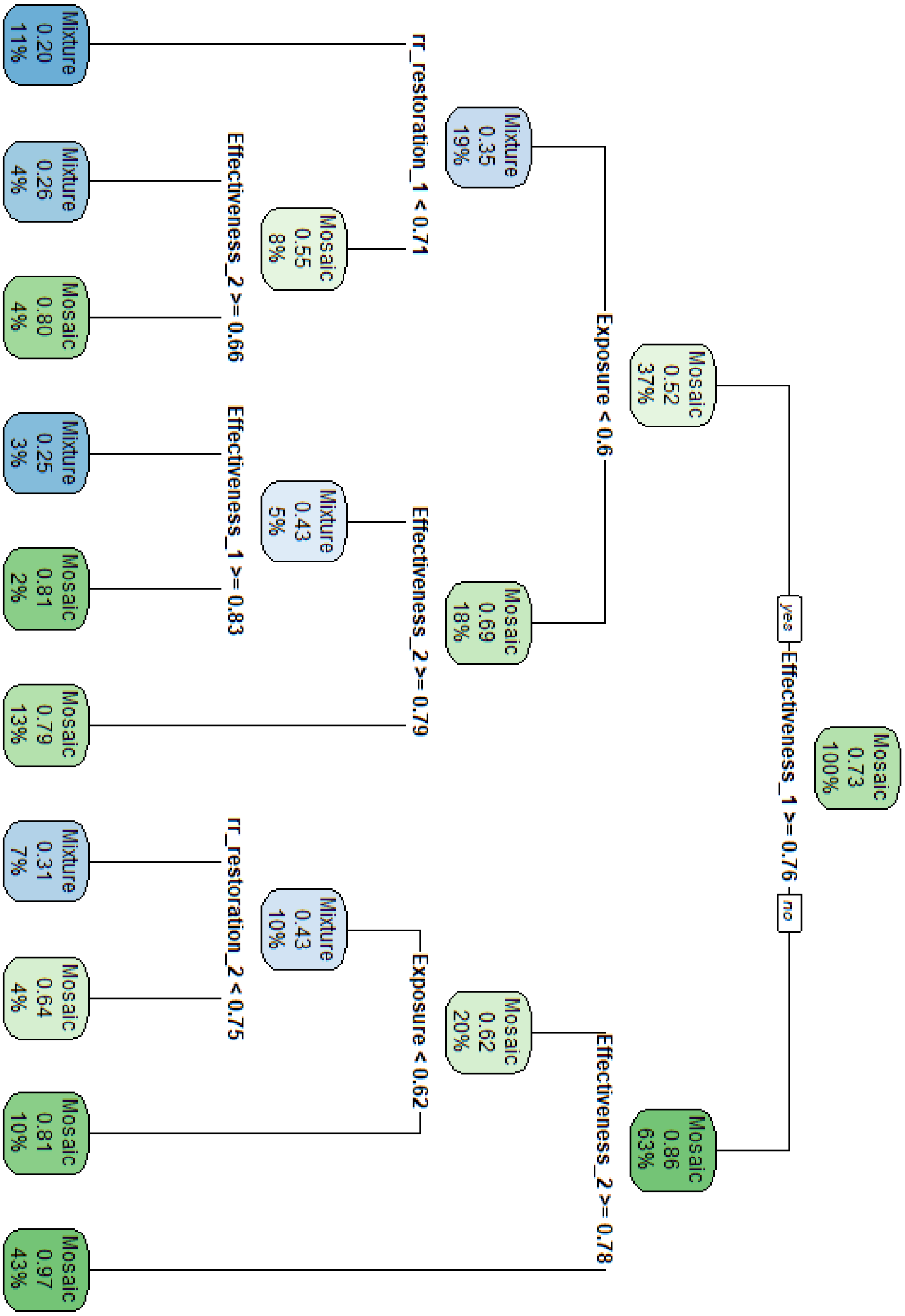

Figure S9. As for Figure 9 of the main text, but analysis based on 250 mosquito generations post-deployment. Classification tree is for a comparison between Panels. Vs micro-mosaics.

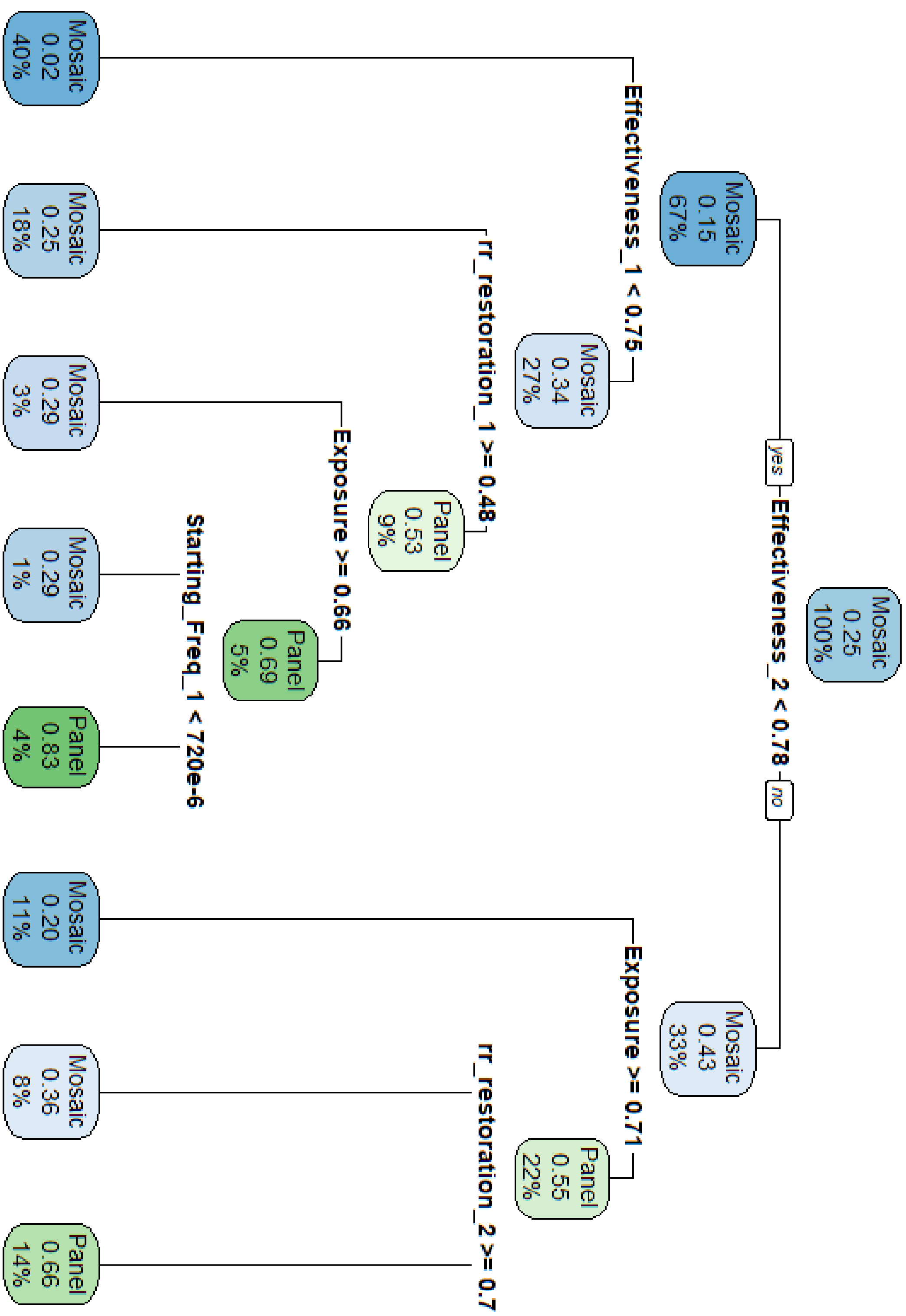
